# Glycerol phosphate acyltransferase 6 controls filamentous pathogen interactions and cell wall properties of the tomato and *Nicotiana benthamiana* leaf epidermis

**DOI:** 10.1101/452102

**Authors:** Stuart Fawke, Thomas A. Torode, Anna Gogleva, Eric A. Fich, Iben Sørensen, Temur Yunusov, Jocelyn K.C. Rose, Sebastian Schornack

## Abstract

The leaf epidermal wall is covered by a cuticle, composed of cutin and waxes, which protects against dehydration and constitutes a barrier against pathogen attack. Cutin monomers are formed by the transfer of 16- or 18-carbon fatty acids to glycerol by glycerol-3-phosphate acyltransferase (GPAT) enzymes, which facilitates their transport to the plant surface. Here we address the dual functionality of pathogen-inducible *Glycerol phosphate acyltransferase 6* (*GPAT6*) in controlling pathogen entry and dehydration in leaves. Silencing of *Nicotiana benthamiana NbGPAT6a* increased leaf susceptibility to the oomycetes *Phytophthora infestans* and *P. palmivora*, whereas stable overexpression of *NbGPAT6a-GFP* rendered leaves more resistant to infection. A loss-of-function mutation of the orthologous gene in tomato (*Solanum lycopersicum*), *SlGPAT6*, similarly resulted in increased susceptibility of leaves to *Phytophthora* infection concomitant with altered intracellular infection structure morphology. Conversely, *Botrytis cinerea* disease symptoms were reduced. Modulation of *GPAT6* expression predominantly altered the outer cell wall of leaf epidermal cells. The impaired cell wall-cuticle continuum of tomato *gpat6-a* mutants resulted in increased water loss and these plants had fewer stomata. Our work highlights a hitherto unknown role for GPAT6-generated cutin monomers in controlling epidermal cell properties that are integral to leaf-microbe interactions and limit dehydration.

## INTRODUCTION

Most epidermal cells of the aerial parts of vascular plants are covered by a hydrophobic extracellular lipid barrier known as the cuticle composed of polymeric cutin and waxes (Yeats & Rose, 2013). The cutin matrix is a highly viscoelastic polymer with low tensile strength (Fich *et al*., 2016) that functions as a transpiration barrier (Schonherr, 1976) and also contributes mechanical strength to the underlying cell wall (Kolattukudy, 1980). Celluloses, hemicelluloses and pectins from the cell wall can be incorporated into the cutin matrix, thereby influencing elasticity and stiffness of the cuticle (López-Casado *et al*., 2007) facilitating expansion and change in composition during plant growth and development and in response to environmental cues (Bargel & Neinhuis, 2005; Underwood, 2012).

Biosynthesis of cutin begins with the esterification of oxygenated fatty acids to glycerol. Cutin monomers contain 16- or 18-carbon fatty acids (Beisson *et al*., 2012) and the glycerol-3-phosphate acyltransferase enzymes (GPAT4, GPAT6 and GPAT8) that transfer these fatty acids to glycerol have specificity for the second carbon of the glycerol (sn-2 position) (Yang *et al*., 2012) as well as phosphatase activity that removes the phosphate group (Yang *et al*., 2010). Mutants of the *Arabidopsis thaliana GPAT4*, *GPAT6* and *GPAT8* genes display reduced levels of C16 and C18 fatty acid monomers (Li *et al*., 2007; Mazurek *et al*., 2016).

*GPAT4* orthologues in *Brassica napus* are highly expressed in the seed coat, periderm and endodermis of roots (Chen *et al*., 2011b) and function in the development of reproductive organs (Chen *et al*., 2014). *GPAT6* is involved in cutin synthesis in *A. thaliana* petals (Li-Beisson *et al*., 2009) and tomato (*Solanum lycopersicum*) fruit (Petit *et al*., 2016) and was found to have multiple functions in stamen development and fertility (Li *et al*., 2012). Analysis of *A. thaliana gpat6* knockout lines demonstrated that GPAT6 is essential for the accumulation of C16 cutin monomers (Li-Beisson *et al*., 2009) and that the enzyme prefers C16 and C18 ω-oxidized acyl-CoA substrates. A glossy fruit mutant of the tomato Micro-Tom cultivar with increased total wax load, but lower levels of total cutin in fruit cuticles, and a much thinner cuticle, (Petit *et al*., 2014) was discovered to be due to a point mutation in the *GPAT6* gene that abolished enzymatic activity (Petit *et al*., 2016). Micro-Tom *gpat6-a* has perturbed pollen formation but is not male sterile (Petit *et al*., 2016).

The cuticle not only controls solute and gas exchange, (Kerstiens, 1996a; Riederer & Schreiber, 2001) but also provides protection against pathogen invasion (Kerstiens, 1996a; Kerstiens, 1996b). *Phytophthora infestans* is an economically important oomycete leaf pathogen of potato (*Solanum tuberosum*) and tomato (Haverkort *et al*., 2008) and can also infect wild tobacco species including *Nicotiana benthamiana* (Becktell *et al*., 2006). *P. infestans* secretes cell wall and cuticle degrading enzymes and forms surface appressoria that support tissue invasion. During early infection stages, *P. infestans* lives as a biotroph, proliferates an extensive intercellular hyphal network within the leaf mesophyll and projects short digit-like haustoria into mesophyll cells to suppress immunity and support infection. At later stages of infection *P. infestans* switches to a necrotrophic lifestyle, killing the host tissue and resulting in necrotic lesions. Other *Phytophthora* species with very similar lifestyles are not restricted to infecting above aerial tissues. For example, the tropical pathogen *P. palmivora* can infect roots and shoots of many vascular and non-vascular host plants (Torres *et al*., 2016). To gain entry into host tissues, pathogens secrete hydrolytic enzymes including cutinases, esterases, lipases and glycanases that destroy the integrity of the cuticle - cell wall continuum (Belbahri *et al*., 2008; Blackman *et al*., 2014). Cutin monomers and cell wall oligosaccharides released during a pathogen attack can serve as damage-associated molecular patterns (DAMPs), allowing the plant cell to mount mitigating defence and wall repair responses, and may also stimulate pathogen colonisation by triggering the formation of appressoria (Gilbert *et al*., 1996). *P. palmivora* forms appressoria when exposed to cutin monomers in vitro (Wang *et al*., 2012) and cutin components have also been reported to trigger spore germination and cutinase expression in the fungus *Botrytis cinerea* (Leroch *et al*., 2013). The ectopic application of cutin monomers during root inoculation with *P. palmivora* was reported to restore full susceptibility to a GPAT mutant (Wang *et al*., 2012) suggesting that the oomycete in part relies on the presence of cutin-derived signals for its invasion process, since reducing GPAT activity enhances resistance to *Phytophthora* infections.

Here we document the importance of GPAT6 in leaf infections by oomycete and fungal pathogens, as well as its contribution to cell wall properties. We found that *GPAT6* transcriptionally responds to *Phytophthora* infection, and that overexpression of *GPAT6* results in increased resistance to oomycete infection. Furthermore, while *gpat6* mutants are more susceptible to *Phytophthora* leaf infection, they display increased leaf resistance to *B. cinerea*, suggesting pathogen lifestyle-specific differences. Changes in pathogen susceptibility are associated with altered thickness of the leaf cell wall plus cuticle, as well as altered transpiration and numbers of stomata. This is reflected in elevated SERK3/BAK1 and ERECTA transcript levels in *gpat6-a* leaves. Cuticle-associated genes are consistently altered in leaves and fruits of *gpat6-a* plants, while more variation exists in genes related to the cell wall and secondary metabolites. While *GPAT6*-like genes have been implicated in flower, fruit and seed development, our work uncovers a new function in leaves of *N. benthamiana* and tomato where *GPAT6* genes influence cell wall and cuticular properties associated with pathogen infection and water regulation.

## RESULTS

### *GPAT6* is induced during *Phytophthora* leaf infections

GPAT enzymes function in root interactions with symbiotic arbuscular mycorrhiza fungi and pathogenic *Phytophthora* oomycetes (Wang *et al*., 2012), but their roles in leaf interactions with pathogens have not been well characterised. To this end, we first identified all GPATs encoded in the tomato and *N. benthamiana* genomes and grouped them based on their phylogenetic relationship to the better characterised *A. thaliana* homologs (Figure S1). This revealed the presence of three *N. benthamiana* genes grouping with *AtGPAT1/2/3* that likely contribute to storage lipid biosynthesis (Zheng *et al*., 2003), and three genes associated with *AtGPAT5/7* that may be involved in suberin biosynthesis (Beisson *et al*., 2007)). The two clades associated with *AtGPAT4/8 and AtGPAT6*, implicated in *A. thaliana* cutin biosynthesis (Li *et al*., 2007; Li-Beisson *et al*., 2009), contain two *N. benthamiana* genes each (Figure S1). An additional group of six *N. benthamiana* genes form a clade together with *MtRAM2* that is distinct from any *A. thaliana GPAT* genes, and were named *NbRAM2A-F*.

Previously published data on *N. benthamiana* root infection by *P. palmivora* reported that *NbGPAT6a*, but none of the other members of the GPAT4/6/8 clade, showed a consistent and significant transcriptional induction during all stages of infection (Figure 1, Suppl. Table 1) (Evangelisti *et al*., 2017).

**Figure 1.**
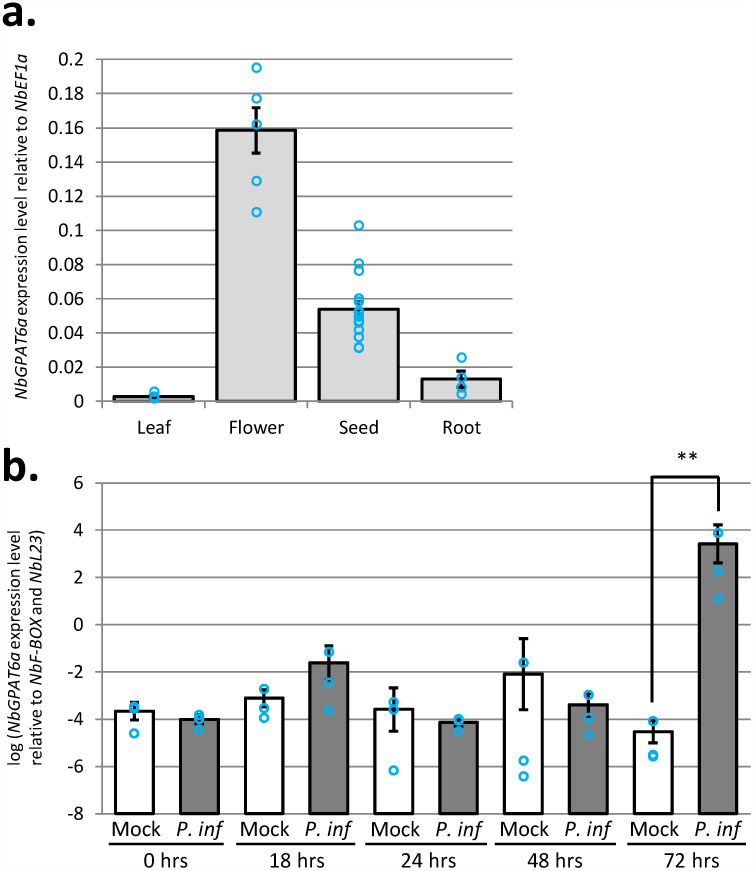
*NbGPAT6a* transcript levels are differently upregulated during *P. palmivora* root colonisation and *P. infestans* leaf colonisation. **a**. Expression level of *NbGPAT6a* in leaf, flower, seed and root tissues. Error bars represent standard error of the mean of at least three biological replicates. **b**. Expression level of *NbGPAT6a* in leaf tissues is significantly increased 72hrs after inoculation with *P. infestans*. Error bars represent standard error of the mean of three biological replicates, ** = *p* < 0.01.

Using quantitative reverse transcription PCR (qRT-PCR) we detected *NbGPAT6a* expression in leaves, flowers, seed and roots, with highest steady state levels in flowers (Figure 1). This is in agreement with *GPAT6* expression patterns in other species (Li-Beisson *et al*., 2009; Li *et al*., 2012; Petit *et al*., 2016).

To test whether *NbGPAT6a* expression levels increase during infection, we infected *N. benthamiana* leaves with *P. infestans* strain 88069 (van West *et al*., 1999) zoospore droplets and measured changes in *NbGPAT6a* expression over time. *NbGPAT6a* was highly induced in leaf tissues at 72 hours post inoculation (hpi) with *P. infestans* (Figure 1). We therefore conclude that *NbGPAT6a* expression is upregulated in roots and leaves infected with *Phytophthora* and that expression levels in leaves are elevated late during infection and so are not part of early, inducible, defence responses.

### Constitutive expression of *NbGPAT6a* renders roots susceptible and leaves resistant to *Phytophthora* infection

To address whether higher *GPAT6a* transcript levels influence *Phytophthora* infection, we generated constitutive overexpression constructs by creating a translational fusion of the genomic *NbGPAT6a* open reading frame (ORF) to the green fluorescent protein (*GFP*) reporter gene under control of the 35S promoter. We first investigated the subcellular distribution of the fusion protein upon transient expression in *N. benthamiana* leaves. GPAT6 is a predicted endoplasmic reticulum (ER)-resident enzyme (Chen *et al*., 2011a) with two transmembrane domains (Figure S2b) and we observed NbGPAT6-GFP signals in the ER of leaf epidermal cells matching the subcellular distribution of ER-targeted red fluorescent protein (RFP) (Figure S2a). We then generated several independent *N. benthamiana* lines that stably and constitutively expressed *NbGPAT6a-GFP* (Figure 2).

**Figure 2.**
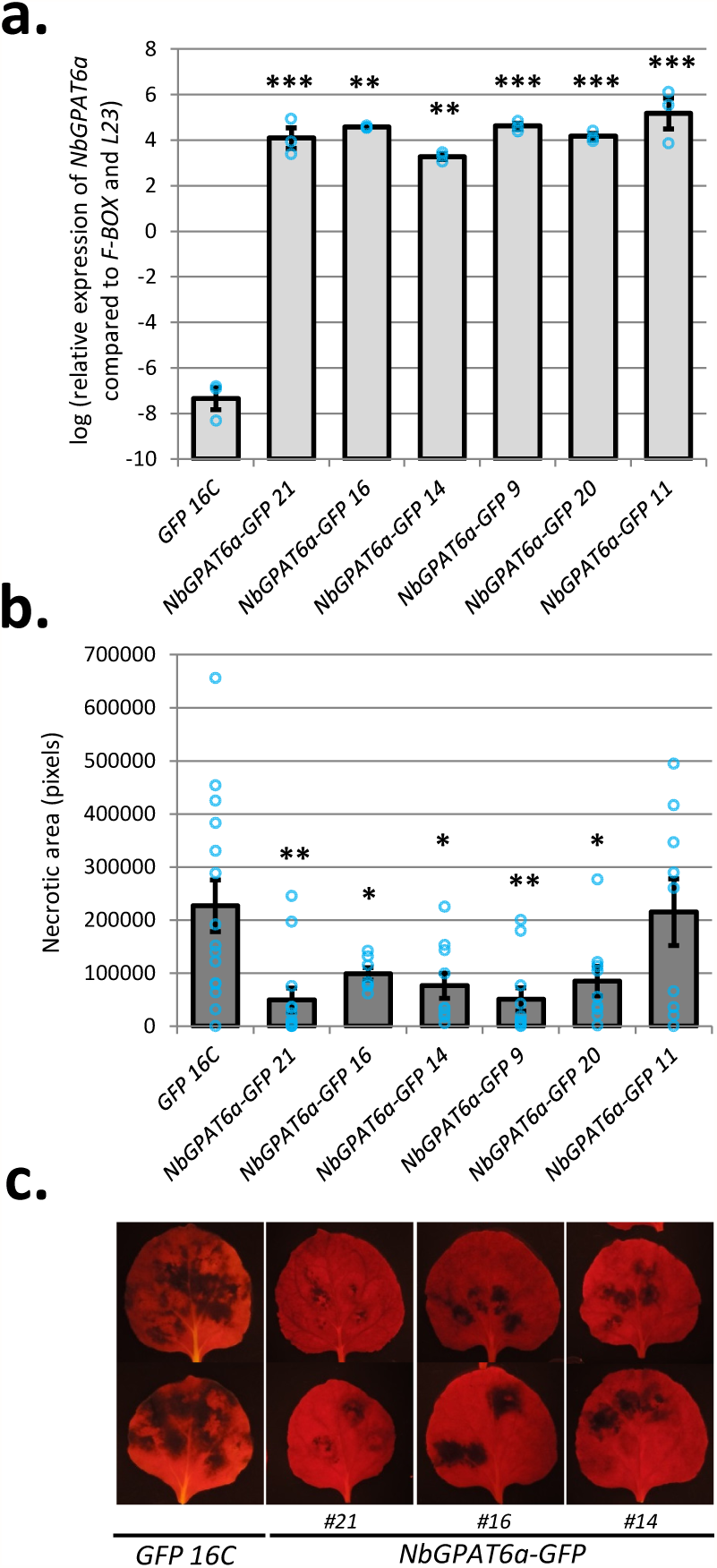
Constitutive overexpression of *NbGPAT6a* renders *N. benthamiana* leaves more resistant and roots more susceptible to infection by *Phytophthora*. **a**. Confirmation of *NbGPAT6a-GFP* overexpression in transgenic *N. benthamiana* leaves, compared to control plants constitutively expressing *GFP* alone, quantified by qRT-PCR. Error bars represent standard error of the mean of three biological replicates. **b**. Quantification of necrotic area present on leaves constitutively expressing *NbGPAT6a-GFP* compared to the control expressing *GFP* alone following inoculation with *P. infestans*. Error bars represent standard error of the mean of at least seven leaves from each transgenic line **c**. Representative images of leaves inoculated with *P. infestans* used to quantify necrotic area.

Overexpression of *NbGPAT6a-GFP* resulted in a 73% increase in total leaf cutin, which was due almost entirely to elevated levels of ω-hydroxyl (OH) fatty acid (-FA) cutin monomers. In particular, levels of hexadecane-dioic acid, ω-hydroxy hexadecanoic acid, ω-hydroxy heptadecanoic acid, ω-hydroxy-octadecanoic acid and 10,16-dihydroxy hexadecanoic acid showed a significant increase relative to GFP16C (control) leaves (Figure S3).

When we tested *NbGPAT6a-GFP* transgenic plants for their resistance to *P. infestans* leaf infections, we found that five of the six lines displayed smaller necrotic areas than the control lines (Figure 2) without affecting overall morphology of *P. infestans* hyphae or haustoria within leaf epidermal cells of two independent *NbGPATa-GFP* transgenic lines (Figure S4). Notably, transient expression of *NbGPAT6a-GFP* in fully expanded leaves followed 24 hrs later by *P. infestans* infection did not alter the extent of disease associated leaf necrosis (Figure S5), suggesting that NbGPAT6-mediated resistance is associated with longer-term leaf development processes.

### Knockdown or knockout of *GPAT6* renders leaves more susceptible to *Phytophthora* infection but more resistant to *Botrytis cinerea*

To test whether reduced *GPAT6* levels cause the opposite phenotype to increased levels, we established a *NbGPAT6a*-specific virus-induced gene silencing (VIGS) construct and demonstrated that it attenuated transcript levels of *NbGPAT6a*, but not those of homologous transcripts (Figure 3 and S6). We found that siNbGPAT6a-mediated VIGS resulted in stronger leaf necrosis upon *P. infestans* infection (Figure 3), suggesting a higher degree of susceptibility.

**Figure 3.**
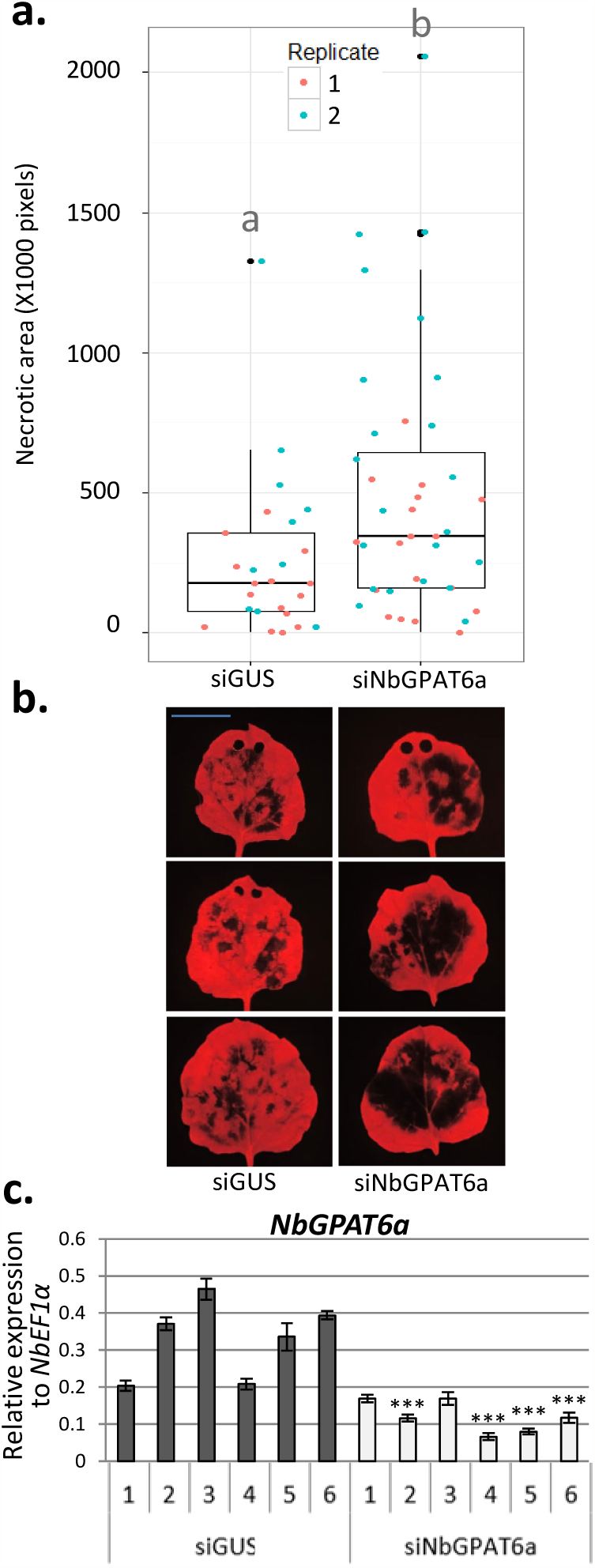
siNbGPAT6a-mediated virus-induced gene silencing results in stronger leaf necrosis upon *P. infestans* infection. **a**. Leaf necrotic area 5 dpi with *P. infestans*, as quantified by lack of red chlorophyll fluorescence, two replicates combined (*p* = 0.0127). Data points for each replicate are denoted by different colours. **b**. Representative UV images of leaf necrosis quantified in a. Scale bar represents 30 mm. Holes at the leaf tip in some images are the result of tissue samples taken for transcript level quantification (these areas were excluded from necrotic area quantification). **c**. Relative expression of *NbGPAT6a* (to *NbEF1α*) following VIGS using siNbGPAT6a or siGUS (control). Error bars represent standard error of the mean of three biological replicates. Student’s t-test used to compare mean relative expression in siNbGPAT6a samples versus lowest of control samples. * = *p* < 0.05 ** = *p* < 0.01 *** = *p* < 0.001.

We next tested whether GPAT6 contributes to *Phytophthora* infection in tomato using a *gpat6-a* mutant in the ‘Micro-Tom’ background (Petit *et al*., 2016). Detached leaf infection assays showed that the tomato *gpat6-a* mutant was more susceptible to *P. infestans* (Figure 4a,b) and *P. palmivora* infection (Figure 4c), as evident by larger lesion sizes and higher expression levels of *P. infestans* sporulation marker transcript levels at 48hrs after inoculation. When investigating infection structures we found that *P. infestans* formed normal, digit-like haustoria (55%) but also singly branched haustoria (45%) in epidermal cells of *gpat6-a* mutant tomato leaves (65 haustoria counted). In contrast, almost exclusively digit-like haustoria (92%, 61 haustoria counted) were formed in wildtype (WT) leaves (Figure S7). Importantly, *gpat6-a* mutants displayed less severe disease symptoms upon infection with the fungal pathogen *B. cinerea* (Figure 4d). Taken together these data demonstrate that attenuating or knocking out the expression of *GPAT6* genes has the opposite effect to *GPAT6* gene overexpression, further supporting an important role for GPAT6 in ensuring full resistance to *Phytophthora* infections.

**Figure 4.**
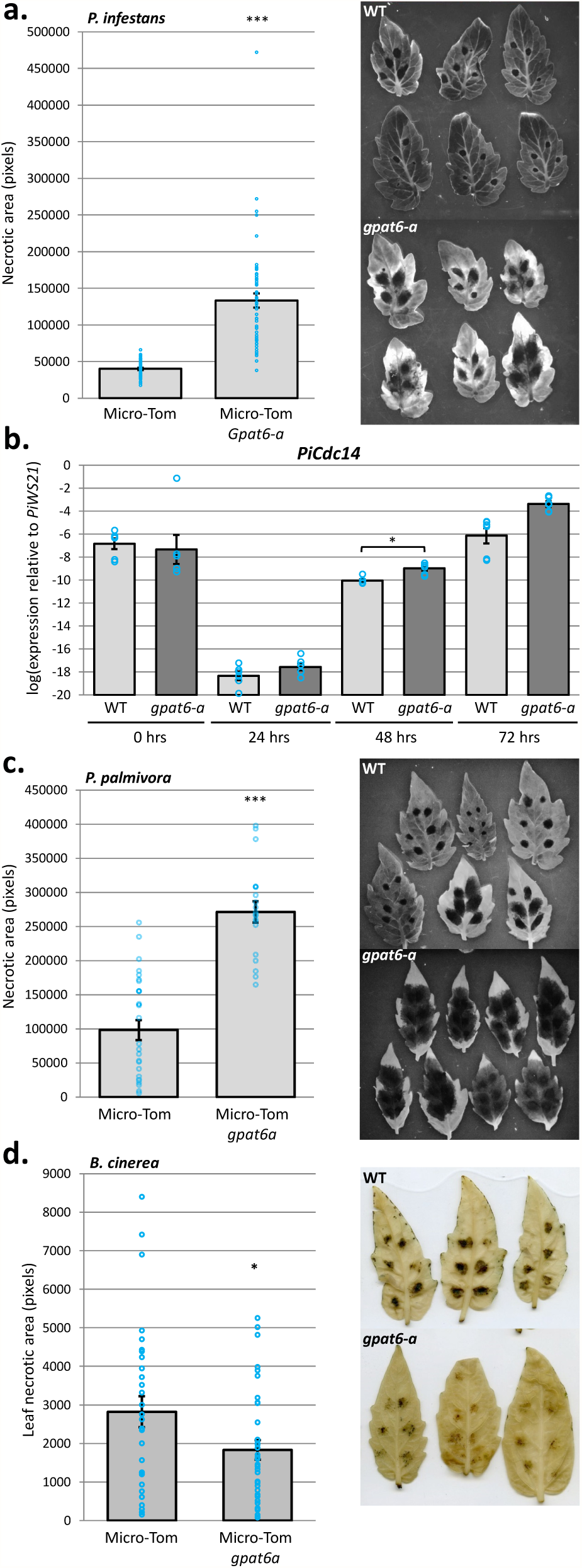
Tomato *gpat6a* mutants are more susceptible to *P. infestans* and *P. palmivora* infection but more resistant to *B. cinerea* infection. **a**. Quantification of Micro-Tom leaf necrotic area 72hrs post inoculation with *P. infestans* for both wild type and *gpat6-a* genotypes. Error bars represent standard error of the mean. Each blue circle indicates the sum necrotic area for one leaf, four inoculation sites. Representative UV images of wild type and *gpat6-a* leaves used for necrotic area quantification are also shown. **b**. Expression level of the sporulation-specific *PiCdc14* (sporulation marker) normalised to *PiWS21* (40S ribosomal protein S3A). Error bars represent standard error of the mean of three biological replicates (six technical replicates indicated by blue circles). **c**. Quantification of Micro-Tom leaf necrotic area 72hrs post inoculation with *P. palmivora* for both wild type and *gpat6-a* genotypes. Errors bars represent standard error of the mean, n = 27 for WT, n = 19 for *gpat6-a*. Each blue circle indicates the sum necrotic area for one leaf, six inoculation sites. **d**. Quantification of leaf necrotic area 6 days post inoculation with *B. cinerea*. Blue circles represent total necrotic area for each leaf (6 droplets of *B. cinerea* spore solution). Error bars represent standard error of the mean, n = 30 for WT, n = 35 for *gpat6-a*. Representative images of leaves used to quantify leaf necrotic area are also shown. This experiment was repeated twice with similar result.

### Modulating *GPAT6* expression alters the thickness of the outer cell walls of the leaf epidermis

The tomato *gpat6-a* mutant was reported to have an altered fruit cuticle structure (Petit *et al*., 2016), which we hypothesised might be associated with the altered *Phytophthora* infection phenotypes described above. We imaged the cell wall and cuticle of *NbGPAT6-GFP N. benthamiana* leaves that displayed different degrees of resistance to *P. infestans* infection, as well as tomato *gpat6-a* mutant leaves, using cryo-scanning electron microscopy (cryo-SEM). We observed that the outer epidermal cell wall was thinner in *NbGPAT6a-GFP* lines compared to those expressing GFP alone (Figure 5a,b), particularly in the line showing the highest *P. infestans* resistance (*NbGPAT6a-GFP* #16).

**Figure 5.**
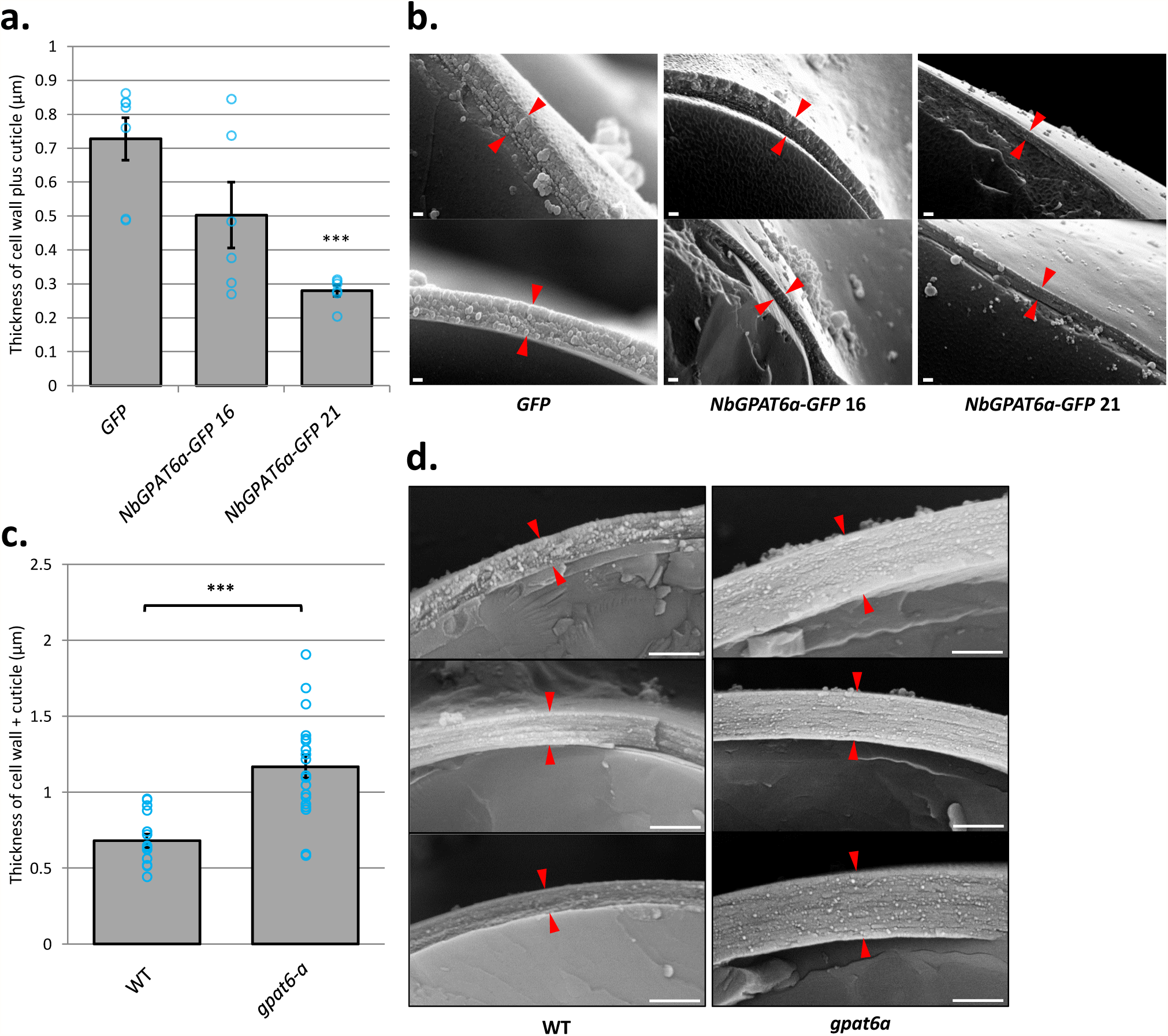
Outer epidermal cell wall thickness is correlated to *GPAT6* expression levels. **a**. Quantification of cell wall plus cuticle thickness in *N. benthamiana* leaves constitutively expressing *NbGPAT6a-GFP*. Blue circles represent mean of 3 measurements per image (n=7 for *GFP*, n=6 for *NbGPAT6a-GFP* 16 and *NbGPAT6a-GFP* 21). Error bars represent standard error of the mean. **b**. Representative cryo-SEM images of transverse fractures used to quantify thickness. Red arrowheads indicate boundary of cell wall plus cuticle. Scale bars represent 200 nm. **c**. Quantification of cell wall plus cuticle thickness in leaves of tomato *gpat6-a* mutants. Blue circles represent mean of 3 measurements per image (n=15 for WT, n=20 for *gpat6-a*). Error bars represent standard error of the mean. **d**. Representative cryo-SEM images of transverse fractures used to quantify thickness. Red arrowheads indicate boundary of cell wall plus cuticle. Scale bars represent 1μm. This experiment was repeated twice with similar result.

Conversely, *gpat6-a* leaf epidermal cells possessed a thicker cell wall (Figure 5c,d & S8). This change in thickness was most prominent in the outer, but not the inner, periclinal wall of both the abaxial and adaxial leaf epidermis (Supp. Fig. S8e). Thus, the cell wall thickness inversely correlated with the level of *GPAT6* expression.

GPAT6 enzymes are known to be involved in cutin biosynthesis (Li-Beisson *et al*., 2009) and *gpat6-a* tomato fruits have increased cuticle permeability to the dye toluidine blue. We found that *gpat6-a* leaves did not show significantly altered permeability when toluidine blue was placed on the upper or lower epidermis while abrasive treatment with bentonite/cellite resulted in full permeability confirming the suitability of our staining procedure (Suppl. Fig. S9). To test for changes in wall porosity we applied Dextran-150kDa-TRITC to WT and *gpat6-a* epidermis. Subsequent fluorescent imaging showed that TRITC labelled Dextran was incorporated to greater extent in the *gpat6-a* mutant, suggesting a larger porosity of the wall (Figure 6).

**Figure 6.**
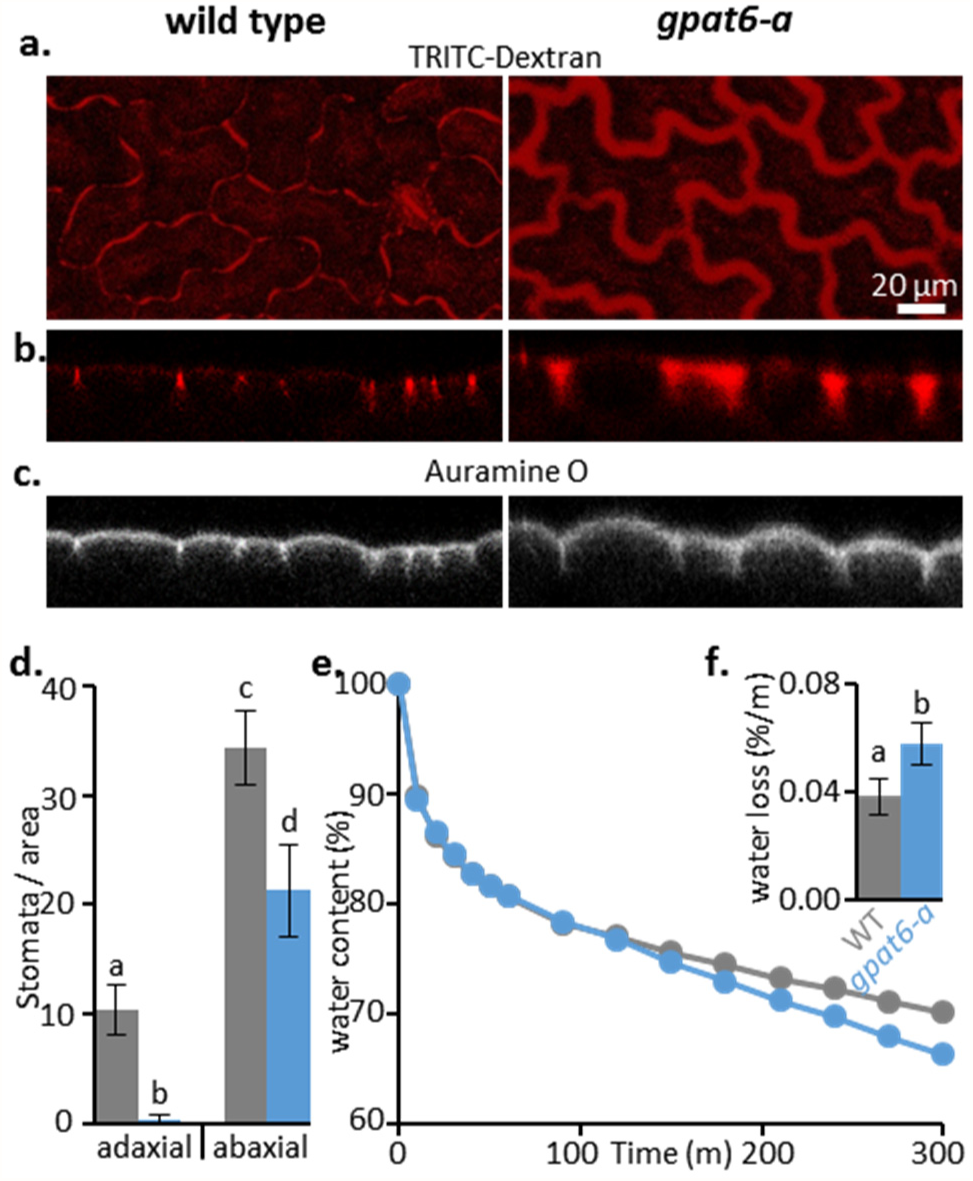
Leaf outer epidermal walls of tomato *gpat6-a* mutant display altered porosity, reduced numbers of stomata and increased susceptibility to desiccation. TRITC-Dextran (150kDa) distribution imaged from above (a) and confocal transects (b) is extended in *gpat6-a* mutant epidermal walls. Auramine O was used to label cutin (c). Stomatal numbers per leaf area are shown in (d); (e,f) water loss over time and total water loss. Error bars represent standard deviation of 6 biological replicates.

Concomitantly, *gpat6-a* tomato leaves showed an increased rate of water loss compared to the WT (Figure 6e,f). This was significant in both the total amount of water loss over time, and the relative water loss over time. We also observed that the *gpat6-a* mutant leaves had fewer stomata than WT (Figure 6d), but that the numbers increased to a level similar to WT when *gpat6-a* plants were grown under high humidity conditions (Suppl. Figure S10). We did not observe changes in stomata numbers or water loss over time in overexpressing *N. benthamiana GPAT6a-GFP* lines with thinner walls (Supp. Figure S11), suggesting that the cuticle permeability was not altered, even though it was thinner. Furthermore, our analysis of the composition and overall architecture of the bulk leaf cell wall using cell wall antibodies did not reveal any significant alterations in *GPAT6-GFP* overexpressing plants or the *gpat6-a* mutant (Supp. Figure S12).

### The *gpat6-a* leaf transcriptome reflects changes in cuticle and cell wall processes and stomatal patterning

To determine the impact of the *gpat6-a* mutation on the tomato transcriptome, we carried out expression analysis of *gpat6-a* and WT leaves from both *P. infestans* infected and control plants (Figure 7), and compared our findings to previous expression data derived from the tomato fruit exocarp (Petit *et al*., 2016).

**Figure 7.**
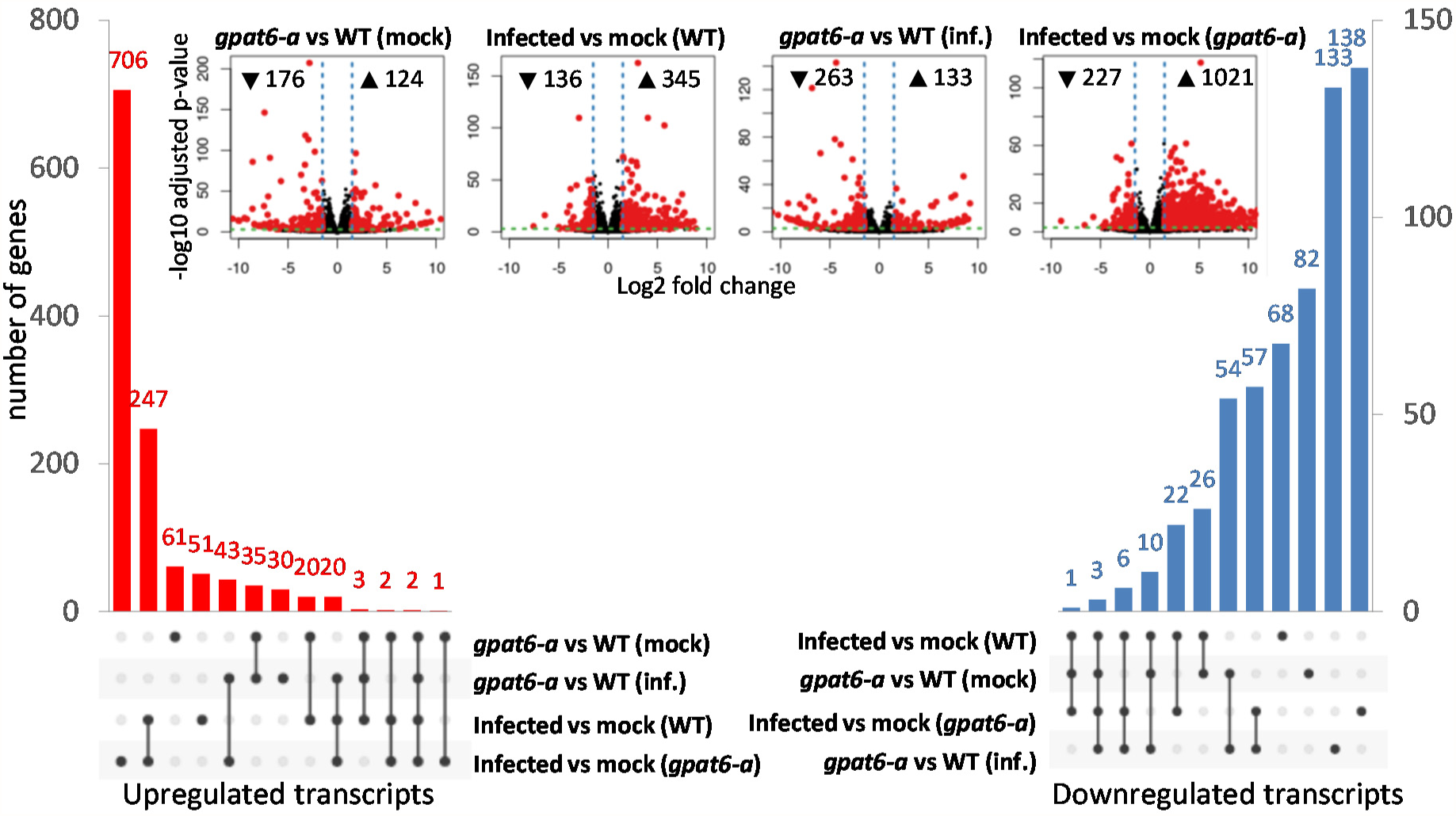
Transcriptome differences in *gpat6-a* vs wildtype tomato cv ‘Micro-Tom’ leaves under control and infection conditions. Water or *P. infestans* 88069 zoospores were applied to detached leaves and harvested 72 hpi. Differentially upregulated (red) and downregulated (blue) transcripts (absolute LFC ≥ 1.5 and adjusted p-value ≤ 10^−3^) and pairwise comparison volcano plots (inset) are shown. This figure is based on Supplementary Dataset 1.

Considering a log fold change (LFC) of 1.5 or higher with an adjusted p-value of <10E-3 significance, we found the expression of 124 genes to be upregulated and 176 to be downregulated compared with uninfected leaves of *gpat6-a* and WT plants (Figure 7, Supplementary Dataset 1). Infection of *gpat6-a* plants resulted in a much higher number of induced genes (1,021) compared to infection in WT leaves (345), congruent with *gpat6-a* leaves being much more susceptible to *P. infestans* infection. Notably, there was a higher proportion of genes associated with immune responses (Pombo *et al*., 2014) induced in *gpat6-a* leaves (Supplementary Dataset 1). The differences in downregulated genes were more moderate, with 227 in *gpat6-a* compared to 136 in WT infected plants.

Petit et al., 2016 highlighted 42 genes associated with lipid, secondary metabolite, and cell wall biosynthesis as differentially expressed in the *gpat6-a* fruit exocarps. We found only 29 of these genes to be altered in the same direction (Suppl. Table 1). Genes belonging to the ‘cuticle’ category largely responded similarly in leaves and fruit exocarp, with 18 of 21 genes similarly differentially expressed in both organs. However, genes in the ‘cell wall’ gene category responded differently and while we found 7 of 10 also repressed in leaves, none of the 5 reported by Petit et al. (2016) was induced. Two genes, annotated as pectin methyl esterase inhibitor and cellulose synthase-like showed opposite expression dynamics between leaves and fruit. Petit et al, 2016 assigned 6 differentially expressed genes to the secondary metabolism category - 3 repressed and 3 induced. Of these, 2 were also repressed and induced in our dataset, respectively and 2 also induced. In summary, the expression levels of cuticle-associated genes were consistently altered in leaves and fruits of *gpat6-a* tomato plants. More variation was observed in the cell wall and secondary metabolite categories. Since alterations in the cell wall were seen to be most extensive in the outer wall of the epidermis, this represents and interesting target for future studies of expression patterns. However, *P. infestans* infection rapidly disrupts epidermal integrity, which represents a major technical challenge.

Even in uninfected leaves, genes associated with defence and immunity such as disease resistance genes, protease inhibitors, pathogenicity-related (PR) genes (*PR-3* and *PR-5*), and chitinases were noted in the group of *gpat6-a* repressed genes and absent from the group of induced genes (Suppl. Dataset 1). Furthermore, seven genes encoding glutaredoxins were repressed in *gpat6-a*, suggesting that this would increase susceptibility to redox stress. Together these data suggest that the increase in susceptibility of *gpat6-a* can, in part, be attributed to constitutively lower levels of defence gene expression in uninfected *gpat6-a* plants.

Six leucine-rich repeat receptor like kinases (RLKs) were induced in *gpat6-a* compared to WT plants (Suppl. Dataset 1) including *SERK3/BAK1* (Solyc01g056655) which showed the strongest level of transcriptional upregulation of all genes differentially expressed between *gpat6-a* and WT leaves, although defence-associated genes were more frequently repressed in *gpat6-a* mutants. A homologue of *ERECTA* homolog (Solyc01g057680) which is associated with negative regulation of stomata density in *A. thaliana* was also induced. The observed induction of both *SERK3/BAK1* and *ERECTA* may be related to the reduced number of stomata in *gpat6-a* leaves. *A. thaliana* homologs of the other induced RLK encoding genes have been associated with pollen tube guidance (Solyc05g025780/LePRK3), (Gui *et al*., 2014) cell wall integrity sensing and resistance to *Fusarium oxysporum* root infections (Solyc01g014147, Solyc01g009930) (Van der Does *et al*., 2017) and regulating cell expansion through the transport of cell wall material (Solyc05g041750/PERK10) (Humphrey *et al*., 2014) but their roles in tomato have yet to be reported.

## DISCUSSION

Previous studies have implicated *GPAT6* in the development of flowers and fruit, but its function in leaves has not been characterised. For example, *A. thaliana GPAT6* is highly expressed in flowers (more than two-fold in petals and sepals compared to other *GPAT* genes) and is known to function in stamen development and fertility (Li *et al*., 2012), while its homologue in tomato has an additional function in fruit cutin biosynthesis (Petit *et al*., 2016). We demonstrate that despite its low steady state expression levels, *GPAT6* fulfils an important role in leaves associated with epidermal outer cell wall properties that confer protection against dehydration as well as infection by *Phytophthora* species.

### Late transcriptional upregulation of *NbGPAT6a* during *P. infestans* infection is a mitigation response

*A. thaliana* GPAT6 is a *sn*2-acyltransferase that is involved in cutin biosynthesis. Our analysis shows that *NbGPAT6a* overexpression raises levels of cutin monomers (Figure S3) indicative of a conserved function. The late transcriptional upregulation of *NbGPAT6* during *P. infestans* infection (Figure 1) can be interpreted either as a pathogen-controlled lipid harvesting strategy or, alternatively, a mitigation response by the plant to tissue damage caused by pathogen colonisation. It was recently hypothesised that obligate biotrophic fungal pathogens may exert a lipid parasitism where the microbe benefits from plant fatty acid production (Jiang, Y. *et al*., 2017; Keymer & Gutjahr, 2018). In this context it was interesting to note a high frequency of aberrantly shaped haustoria in infected *gpat6-a* tomato (Figure S4). *P. infestans* haustoria are intracellular structures and characteristically digit-shaped, although infrequently branched haustoria can also be observed (Blackwell, 1953). Whether an altered lipid metabolism in *gpat6-a* tomato leaf epidermal cells may impact on their development requires additional confirmation using other lipid biosynthesis mutants in tomato. Nevertheless, *gpat6-a* tomato mutants were more susceptible to both *P. infestans* and *P. palmivora* (Figure 4) suggesting that the observed alteration in haustorium morphology does not impair infection and that the oomycete does not extensively rely on cutin monomers provided through GPAT6. It may also be interesting to infect *gpat6-a* tomato with obligate fungal pathogens such as *Oidium neolycopersici* to address whether lipid parasitism is linked to an obligate biotrophic lifestyle. A *Phytophthora*-controlled lipid harvesting scenario is also unlikely because overproduction of cutin monomers through overexpression of *NbGPAT6a-GFP* did not result in increased pathogen-caused symptoms, indeed the leaves were more resistant to *P. infestans* infection (Figure 2). Furthermore, *Phytophthora* cuticle and cell wall-degrading enzymes likely release cutin monomers into the apoplast to enable sufficient uptake by the oomycete for continued infection in a WT situation. We therefore propose that late transcriptional upregulation of *NbGPAT6a* during *P. infestans* infection is a mitigation response to tissue damage. Alternatively, it may be part of a delayed defence response as various hydroxy fatty acid compounds have been implicated in disease resistance (Schweizer *et al*., 1996; Hou & Forman Iii, 2000; Wang *et al*., 2000) including against *P. infestans*.

### Differences in oomycete and fungal leaf infections may be attributable to their lifestyles

A range of mutations have been reported to both increase leaf cuticle permeability and increase pathogen susceptibility (Tang *et al*., 2007). However, these mutants are all more resistant to *B. cinerea* (Ziv *et al*., 2018). Explanations for this apparent paradox include release of disease resistance activators, antifungal diffusible components and improved uptake of elicitors (Ziv *et al*., 2018) (Tang *et al*., 2007). A key factor in the ability of pathogens to infect these mutants may be their infection biology or lifestyle. *Phytophthora* pathogens are hemibiotrophs which initially require living host cells for infection (Fawke *et al*., 2015), whereas *B. cinerea* is considered a necrotroph which immediately kills the tissue (Van Kan *et al*., 2017). This may explain why *gpat6-a* tomato leaves display resistance to *B. cinerea*. While a reliance on plant lipid biosynthesis was recently demonstrated for an obligate biotrophic fungal pathogen (Jiang, Yina *et al*., 2017), this remains to be reported for oomycetes.

### The leaf cuticle layer may impose a physical restraint upon the outer facing epidermal cell wall

Our results suggest that GPAT6 influences the physiological properties of the cell wall. We show that loss of *GPAT6* increased the thickness of the wall (Figure 5c,d & S8) and that overexpression of *NbGPAT6a-GFP* led to higher levels of cutin monomers, reducing the thickness of the wall (Figure 5a,b). Interestingly, this effect was mainly observed at the epidermal walls facing the outside environment, which are also the only walls lined by a cuticle (Figure S8e), while the overall composition of the leaf bulk cell wall was not altered (Supp. Figure S12). This suggests that the outer epidermal cell wall may respond to its cutin status and adapt its thickness, possibly through mechanical or biochemical sensing. An instantaneous and reversible increase in thickness of the cell wall has been observed previously upon abrasion of the cuticle (Xia *et al*., 2009). This suggests that physical properties of the wall allow it to flexibly expand in diameter and that cutin monomers contribute to preventing excessive expansion. Similar increases in wall thickness were reported when a tomato cutin synthase was mutated leading to a substantially thinner fruit cuticle in tomato (Yeats *et al*., 2012). Although we observed that *gpat6-a* mutant leaves have a thicker cell wall and cuticle while those of leaves constitutively expressing *NbGPAT6a* are thinner, there was not such a clear contrast in terms of the rate of water loss or stomata numbers. Specifically, *gpat6-a* leaves loose more water over time than WT (Figure 6e,f) whereas leaves constitutively expressing *NbGPAT6a* loose the same amount of water as WT (Figure S11). This suggests that a lack of cutin monomers during development has a significant impact on permeability, whereas an excess of cutin monomers can be tolerated or compensated for by the plant and has no effect on permeability.

The increased porosity of *gpat6-a* mutants with thicker walls may result from the looser packing of wall components or from the low levels of cutin monomers in the wall. This in turn may cause the observed increased rate of water loss, which is compensated for by altering the number of stomata in non-water saturated atmosphere.

## MATERIALS AND METHODS

### Statistical analysis

Levene’s tests were applied to check for heteroscedasticity between treatment groups. Following this the appropriate two-sample t-test was applied, accounting for equal or unequal variances, to assess whether the means of two different treatment groups were significantly different, based on alpha = 0.5. Figures are labelled with asterisks to indicate p value range, i.e. * = p≤0.05, ** = p≤0.01, *** = p≤0.001

### Microbial strains and cultivation

*P. infestans* strain 88069, previously described in (van West *et al*., 1998), was grown at 18°C in the dark on rye sucrose agar plates. Zoospores were harvested from 14-day old plates by adding 6ml cold sterile H_2_0, incubating in the dark at 4°C for 45mins - 1 hour then in the light at room temperature for 30 mins and extracting the liquid by pipetting. Approximately 30spores/μl sterile H_2_0 was then used immediately for infection assays.

*P. palmivora* strain P16830-YKDEL was previously described (Rey *et al*., 2013). *P. palmivora* was grown in a Conviron A1000 Reach-In Plant Growth Chamber at 25°C, 700μmol intensity. For subculturing, rye sucrose agar plates were used with the addition of 50μg/ml G418 (geneticin) to select for transformants. For production of zoospores, agar plates containing 10% unclarified V8 vegetable juice were used with the addition of 50μg/ml G418 (geneticin). Harvesting of zoospores was performed as for *P. infestans* described above.

*Botrytis cinerea* R190/11/3, isolated from *Geranium* by Robert Saville in 2011 (NIAB-EMR, East Malling, UK) was grown on potato dextrose agar plates in a Conviron A1000 Reach-In Plant Growth Chamber at 25°C, 700μmol intensity and subcultured by excising an agar plug containing conidiophores and inverting it onto a fresh plate. Conidia for infection assays were harvested from 7-day old potato dextrose agar plates but adding 6ml cold sterile H_2_O, incubating in the light at room temperature for 1 hour then gently agitating the conidiophores with a spatula to release the conidia. The concentration was adjusted to approximately 30 conidia/μl sterile H_2_O.

### Leaf infection assays

10μl droplets of zoospore or conidia suspension were placed onto the abaxial side of the leaf between the veins. Inoculated leaves were then incubated in a humidified chamber for the stated time before imaging. UV images of *P. infestans* or *P. palmivora* lesions were obtained using a blue light table and digital camera set to long exposure.

### Phylogenetic analysis

Protein sequences from *N. benthamiana*, *M. truncatula* and *S. lycopersicum* homologous to *A. thaliana* GPATs were identified by BLASTp search of the NCBI database (https://blast.ncbi.nlm.nih.gov/Blast.cgi) using AtGPATs 1-9 as queries (sequence accession numbers are listed in Figure S1). Obtained sequences were then aligned using MUSCLE (http://www.ebi.ac.uk/Tools/msa/muscle/) and a phylogenetic tree constructed using PhyML (http://atgc.lirmm.fr/phyml/) with the Phylogeny.fr web tool (http://www.phylogeny.fr/). Branches with <50% bootstrap support (100 iterations) were collapsed. The tree presented in Figure S1 was rendered using TreeDyn (http://www.treedyn.org/) and annotated using GIMP (https://www.gimp.org/).

**Confocal microscopy** was performed using a Leica SP8 equipped with a white light laser. GFP excitation wavelength: 488nm, Tdtomato excitation wavelength: 554nm.

### qRT-PCR

Normalisation of Cp values to an internal control was performed against *NbEF1a*, *NbF-BOX* or *NbL23* (Liu *et al*., 2012) for quantification of *N. benthamiana* transcripts and against *PiWS21* (Yan & Liou, 2006) for *P. infestans* transcripts.

### Expression analysis

Leaves of 6-week old tomato cv ‘Micro-Tom’ or *gpat6-a* mutant plants were subjected to a detached leaf infection assay (see above) and either zoospore suspension or water were applied to the lower epidermis. Leaf discs were harvested 72hrs after inoculation. Three biological replicates per sample were obtained and subjected to RNA extraction and poly(A) selection. cDNA library preparation was performed with the TruSeq^®^ RNA Sample Preparation Kit (Illumina, US) according to the manufacturer’s protocol. cDNA sequencing of the 12 samples was performed with Illumina NextSeq 2500 in 100 paired-end mode (Genewiz). Raw reads were subjected to quality control with FastQC (https://www.bioinformatics.babraham.ac.uk/projects/fastqc/) and then aligned back to the *S. lycopersicum* reference genome ITAG3.1 (ftp://ftp.solgenomics.net/tomato_genome/annotation/ITAG3.1_release/) using STAR (version 2.5.2b) aligner. Raw counts were obtained with FeatureCounts (Liao et al. 2014), and only uniquely mapped and properly paired reads were considered further. Differentially expressed genes were identified with the DESeq2 Bioconductor package (Love et al. 2014) following four pair-wise comparisons. Differentially expressed genes (absolute LFC ≥ 1.5 and adjusted p-value ≤ 10-3) were used to generate volcano plots and upset plots using upsetR package (Conway *et al*., 2017).

#### Cryo-scanning electron microscopy

Cryo-SEM was performed on 6 week old *N. benthamiana* and tomato leaves using a Zeiss EVO HD15 with a Quorum cryo-prep deck and cryo-stage. Leaf sections were mounted, frozen, positioned inside the cryo-prep deck and then fractured using a blade to allow for cross sectional imaging. Sublimation of samples for 3 mins was used to remove surface ice and a 5nm platinum coating was applied prior to imaging via secondary electron detection.

### Cell wall porosity analysis

Disks (5mm diameter) were excised from leaves using a cork borer and incubated for 1h at room temp with TRITC:Dextran (0.1 mg/ml, 150k mw, TDB Consultancy AB) and Auramine O (0.01% w/v, Sigma). Images were acquired using a Leica SP8 (excitation wavelengths: TRITC 561; AuramineO, 458).

### Water loss/dehydration analysis

Whole leaves were harvested, placed in a ventilated oven (30°C; MAXQ-6000, Thermos Scientific) and weighed over a timecourse.

## ACKNOWLEDGEMENTS

The authors would like to thank Clement Quan, Edouard Evangelisti, Liron Shenav, Firas Bou Daher and Ray Wightman (all SLCU, Cambridge) for technical and training support, Robert Saville (NIAB-EMR, East Malling) for providing *Botrytis cinerea*, Sam Brockington (Plant Sciences, Cambridge) for advice on phylogenetic analysis, Paul Knox (University of Leeds) for providing monoclonal antibodies and Christophe Rothan (INRA, Paris) for providing tomato seeds.

## SUPPLEMENTARY METHODS

### Transient overexpression of *NbGPAT6a-GFP* in *N. benthamiana* leaves

The *NbGPAT6a* ORF (SolGenomics *N. benthamiana* Genome v1.0.1 Gene ID: Niben101Scf25069g01006.1) was amplified from *N. benthamiana* genomic DNA. It was first cloned into the Gateway entry vector pENTR/D-TOPO and subsequently recombined into pK7FWG2 (Karimi *et al*., 2013), transformed into competent *Agrobacterium tumefaciens* strain GV3101 cells via electroporation (1500V, 5ms) and grown on LB containing rifampicin/gentamycin/spectinomycin (all 50μg/ml) at 28°C. Overnight grown liquid cultures were centrifuged at 4000rpm for 10 mins and the bacterial pellet resuspended in a solution of 10mM MES and 10mM MgCl2 to an OD600 of 1.0. Acetosyringone was then added at 200μM concentration and the suspension incubated under shaking for a further 2-3 hours at 28°C. Finally, the suspension was infiltrated into the abaxial side of 4 week old *N. benthamiana* leaves using a syringe barrel without needle. For leaf infection assays, leaves were detached 24hrs post-infiltration for immediate inoculation with zoospore or conidiophore suspensions. When performing microscopy to study subcellular localisation of NbGPAT6a-GFP leaves were detached 4 days post infiltration.

### Toluidine Blue treatment

Leaves were treated with a mixture of Bentonite and Celite (0.02% and 1% w/v, respectively) using cotton wool swabs, or treated with water alone as a control. Leaves were then stained with droplets of Toluidine Blue (TBO, 30 μl, 0.1%) for an hour at room temperature, washed with water and then imaged on a flatbed scanner.

### Cutin analysis

One mature leaf from each of three individual 5.5-week old *N. benthamiana* plants expressing GFP16C (control) or NbGPAT6a-GFP was harvested. Images of the leaves generated with a flatbed laser scanner (Epson V30) were analyzed using ImageJ (https://imagej.net) to determine the leaf surface area in order to calculate the amount of cutin per cm^2^ leaf area. The leaf samples were then subjected to delipidation using solvents containing 0.01% BHT (butylated hydroxytoluene). For each sample, 30 mL of isopropanol was preheated in a glass vial to 85°C and leaf samples were cut into squares (~0.5 cm^2^), added to the hot isopropanol, and incubated at 85°C for 20 minutes. The vials were then cooled to room temperature and agitated at 250 rpm for 1 hour, then the supernatant was poured off. The tissue was then sequentially washed with each of the following solvents: isopropanol (2nd wash), 1:2 methanol: chloroform, 2:1 methanol: chloroform, methanol. A total of 30 mL of solvent was used for each wash, with agitation for at least 2 hours and up to overnight (2:1 methanol: chloroform step). After the final wash, the tissue was dried under a stream of nitrogen gas and then placed in a vacuum desiccator overnight. Cutin depolymerisation was performed as described (Li-Beisson *et al*., 2013), except that twice the amount of the reaction medium was used and the 60°C incubation step was performed overnight. The final products were filtered through filter paper (VWR #28333-021) into clean vials. An 1 mL aliquot of each cutin sample was dried by heating at 40°C under a stream of nitrogen, then derivatised with 50 μL each of 1 μg/ul pyridine and BSTFA (N,O-bis(trimethylsilyl)trifluoroacetamide) for 10 minutes at 90°C. The samples were dried again by heating to 50°C under nitrogen and re-suspended in 100 μL of chloroform. The samples were then analysed by gas chromatography (GC) as described (Bolger *et al*., 2014). Cutin levels were normalised based on the internal standards and the surface area of the leaves (taking into account abaxial and adaxial surfaces).

### Cell-wall component enzyme linked immune sorbent assay (ELISA)

Leaf material was harvested, frozen, finely ground, and processed into an alcohol insoluble residues (AIR) via sequential washes with 80%, 90%, and 100% (v/v) ethanol, acetone, and finally methanol/chloroform (2:3 v/v) and left to dry overnight. AIR (5 mg) was extracted sequentially with 50 mM 1,2-cyclohexanediaminetetraacetic acid (CDTA), and 4 M KOH. CDTA and KOH samples were diluted 1 in 10 into PBS (KOH samples were neutralised first with glacial acetic acid) and used to coat microtiter plates (maxisorb, Nunc) and ELISAs were performed as described (Torode *et al*., 2016) using cell wall specific monoclonal antibodies(Cornuault *et al*., 2017)

### Prediction of transmembrane domains

used TMHMM Server v. 2.0 (Sonnhammer *et al*., 1998)

### Data availability statement

The raw fastq data from the expression analysis are accessible at http://www.ncbi.nlm.nih.gov/sra/ with accession number SRP158564. This data forms the basis of Figure 7 and Suppl. Dataset 1. There are no restrictions on data availability.

## AUTHOR CONTRIBUTIONS

SS, TA, AG, SF, JR designed the research; SF, TA, AG, EF, IS, TY, SS performed research; SF, TA, AG, EF, IS, JR, SS analysed data; SF, JR and SS wrote the paper.

